# Integrated ACMG approved genes and ICD codes for the translational research and precision medicine

**DOI:** 10.1101/2023.01.14.524076

**Authors:** Raghunandan Wable, Achuth Suresh Nair, Anirudh Pappu, Widnie Pierre-Louis, Habiba Abdelhalim, Khushbu Patel, Dinesh Mendhe, Shreyas Bolla, Sahil Mittal, Zeeshan Ahmed

## Abstract

Timely understanding of biological secrets of complex diseases will ultimately benefit millions of individuals by reducing the high risks for mortality and improving the quality of life with personalized diagnoses and treatments. Due to the advancements in sequencing technologies and reduced cost, genomics data is developing at an unmatched pace and levels to foster translational research and precision medicine. Over ten million genomics datasets have been produced and publicly shared in the year 2022. Diverse and high-volume genomics and clinical data have the potential to broaden the scope of biological discoveries and insights by extracting, analyzing, and interpreting the hidden information. However, the current and still unresolved challenges include the integration of genomic profiles of the patients with their medical records. The disease definition in genomics medicine is simplified, when in the clinical world, diseases are classified, identified, and adopted with their International Classification of Diseases (ICD) codes, which are maintained by the World Health Organization (WHO). Several biological databases have been produced, which includes information about human genes and related diseases. However, still, there is no database exists, which can precisely link clinical codes with relevant genes and variants to support genomic and clinical data integration for clinical and translation medicine. In this project, we are focused on the development of an annotated gene-disease-code database, which is accessible through an online, cross-platform, and user-friendly application i.e., PAS-GDC. However, our scope is limited to the integration of ICD-9 and ICD-10 codes with the list of genes approved by the American College of Medical Genetics and Genomics (ACMG). Results include over seventeen thousand diseases and four thousand ICD codes, and over eleven thousand gene-disease-code combinations.

## 1. Introduction

Symptom-driven medicine has become the domain of medical research for the past decade [1, 2]. However, some challenges arise when focusing on the symptoms rather than the disease. Patients with life-threatening diseases might not feel pain and seek professional help. Thus, personalized treatment to help manage and identify those patients, with the help of precision medicine is needed to effectively diagnose and provide the most optimal actions needed for a patient. [3, 4, 5]. Precision medicine is a multi-disciplinary field that utilizes the clinical and multi-omics data of an individual to create patient-specific treatment plans and diagnoses [4, 7, 8]. Clinical data is most familiar to clinicians and patients as the medium that communicates personal and health information between the provider and patient. Genomic information is stored within various databases that include but are not limited to ClinVar, CNVD, Cochrane Library, Disease Ontology, Disease Enhancer that allow for gene annotation [4]. However, there is a lack of standardized, comprehensive databases that consolidate known gene-disease relationships. Furthermore, there is no known database that connects International Classifications of Disease (ICD), mediated by the World Health Organization (WHO), with the list of 73 genes compiled by the American College of Medical Genetics and Genomics (ACMG), whose mutations are known to be causative of disorders and disease [9].

The evolution from the first use of the word gene to our current understanding has launched a new scientific age. On an introductory level, the chemical structure of the genome is in the form of deoxyribose nucleic acid (DNA) which is comprised of a double helix with a pair of nucleotides connected through a hydrogen bond [1, 10, 11]. These alternating patterns of nucleotides (adenine, cytosine, guanine, and thymine) encode the instructions for all the proteins in our body, yet only a fraction of the entire genome contains protein coding sequences [6, 12]. The goal of genomic medicine is to isolate and examine the mutations in these sequences that lead to diseases [6, 13, 14]. This objective is observable in the link between sickle cell anemia and the mutation in protein encoding the hemoglobin once the genome is sequenced [1]. The sequencing and understanding of these mutations have been made possible by Next-Generation Sequencing (NGS) [15]. Currently, Illumina sequencing is the most popular sequencing technology due to its accuracy, cost, and speed [16]. Illumina sequencing belongs to a family of NGS technology that produces short reads (50–300 base pairs), with the most notable other technology in this category being Ion Torrent sequencing [17]. After the sequencing data is collected, it is displayed and shared as a FASTQ file. Each sequence stored in the FASTQ file has four corresponding lines of text. These lines contain information such as the sequence identifier, nucleotide sequence, a “+” sign to indicate the end of the sequence, and a line of quality values reported in the American Standard Code for Information Interchange (ASCII) characters [6, 18]. Using gene information in a FASTQ file, algorithms map the reads to the reference genome and stores it in a Sequence Alignment Map (SAM) or its binary equivalent (BAM) file [19]. From the SAM file, variant call format (VCF) files are created which store information regarding variations, insertions, and deletions. [6]. Whole Genome Sequencing (WGS) and Whole Exome Sequencing (WES) are two types of NGS which are more accurate methods of DNA sequencing and are used to find variants in a DNA sequence [20]. While WGS sequences the whole genome, WES sequences only the protein-coding sections [21].

Recent developments in sequencing technologies have greatly aided in long read sequencing and integration of genomic data. However, challenges arise when integrating heterogenous data such as clinical and genomic data. Electronic health records (EHR) contain large volume of data that cannot be processed at a fast and efficient rate on local servers. Thus, it is vital to use high-performance computing to process this data [22]. We recently created a Java-based Whole Genome/Exome Sequence Data Processing Pipeline (JWES), a free, open-source pipeline that processes WES/WGS as well as EHR data, stores information, and provides user-friendly visual analysis [23]. Information is parsed through and analyzed by JWES’s connection to a high-performance computing cluster allowing for efficient analysis of big data [23]. Due to the personal nature of the data included in EHRs, it is imperative that safeguards are placed to protect the confidentiality of such data [24]. In the genetics field, ACMG is a medical organization that is responsible for guidelines internationally accepted for variant interpretation along with improving health through genomics and medical genetics [9]. ACMG is responsible for publishing and providing recommendations for clinical exome and genome sequencing that provides a universally accepted platform for scientists to work and discover any new incidental findings [9]. Presently, 73 genes have been proposed by ACMG which are known to be of importance to disorders and can be clinically acted on by an accepted way of intervention [25]. These genes provide significant medical value as it allows for improved clinical treatment [9, 25].

The duality of information stored by genomic and clinical data in a single network would form a comprehensive patient profile that creates the possibility for individualized health care. However, there is no system that integrates the two data types and standardizes the data according to international academic standards [1, 23, 26]. This shortcoming allows symptom-based treatments to be normalized as the default approach to patient care, and to challenge the standard model a solid connection must be made between clinical and genomic data [23, 26]. Even with the latest sequencing technologies, the format and robustness of raw DNA and RNA files, especially WES, are not well suited for current EHR systems [27]. Raw genomic files must undergo various processing procedures before being able to be visualized and used by non-bioinformaticians [23]. Combined with the intense computing environment needs for maintaining an EHR system [22], there is an infrastructural component to also consider. However, recent developments in the field highlight some promising outcomes in the creation of a unified genomic-EHR system. PROMIS-APP-SUITE (PAS)-Gen mobile application is a publicly available iOS app that leverages a database of over 59,000 coding and non-coding genes along with 90,000 gene disease associations [20]. It was created with the intention of assisting academic researchers and medical professionals in understanding the dynamic between disease and genes [20]. This interface shines a light on the future integrated databases could have with some restructuring.

The organization of healthcare information is largely based on a label-based systems. On a global scale the WHO created the standardized ICD codes while the Food and Drug Administration (FDA) maintains the National Drug Code (NDC) [28]. NDC serves as an identifier for prescription and over the counter drugs as well as insulin. This database contains information pertinent to the commercial sale of drugs, including manufacturer and packing details [29]. In this project, we have designed and implemented a relational database and interactive online web application that connects genomic and clinical data, allowing for a user to discover gene and disease relations along with their respective ICD codes. We hypothesize that our web application can assist healthcare providers and clinicians to create a more personalized treatment approach by observing gene-disease-ICD. However, the scope of research is limited to only ACMG genes.

## 2. Material and Methods

Our methodology was divided into three main sections. Firstly, we were focused on curating and integrating the genomic and clinical data. Then, we focused on designing and modeling a new relational database to facilitate data manipulation. The last step highlights the implementation of our efficient and user-friendly online web application PAS – Gene Disease Code (GDC) to facilitate integrated search of clinical and genomic data all in one place.

### 2.1. Gene-disease-code data curation and integration

PAS-GDC website uses the seventy-three approved genes from ACMG as well as the ICD codes to curate the data that powers the search engine (Table 1). ICD-9 and ICD-10 codes were utilized in the creation of our relational database. While the ICD-9 codes were proposed by the WHO to provide a unified system to present mortality statistics, the ICD-10 codes were implemented for inpatient procedures in hospitals [28]. Additionally, the structure of ICD-9 and ICD-10 codes are vastly different where ICD-9 codes are numeric, and ICD-10 are alphanumeric. The PAS-GDC website currently holds 2,101 ICD-9 codes and 2,589 ICD-10 codes that were manually curated for search functionalities. Additionally, there are 7,918 and 11,799 gene-disease combinations for ICD 9 and 10 codes, respectively (Table 2). Two Excel Sheets were curated containing up-to-date information regarding each of the seventy-three actionable genes, their relevant diseases, and relevant ICD-9 and 10 codes. For easy translation from the excel sheet to Structured Query Language (SQL) relation, a Python extraction, transfer, and load script was written. Running this script provides the user with a text file containing the genes, diseases, and ICD codes, which can be copied and pasted into SQL to create two relations containing all information from both Excel Sheets (Figure 1).

**Table 1.** List of American College of Medical Genetics and Genomics (ACMG) genes. Table 1 includes a list of 73 ACMG genes for which specific mutations are known to be causative of disorders with defined phenotypes that are clinically actionable by an accepted intervention. The disease phenotype associated with each gene is also included.

| Number | Genes | Name of the Disease |
| --- | --- | --- |
| 1 | BRCA1 | Hereditary breast and ovarian cancer |
| 2 | BRCA2 |  |
| 3 | PALB2 |  |
| 4 | TP53 | Li-Fraumeni syndrome |
| 5 | STK11 | Peutz-Jeghers syndrome |
| 6 | MLH1 | Lynch syndrome |
| 7 | MSH2 |  |
| 8 | MSH6 |  |
| 9 | PMS2 |  |
| 10 | APC | Familial adenomatous polyposis |
| 11 | MUTYH | MYH-associated polyposis; adenomas, multiple colorectal, FAP type 2; colorectal adenomatous polyposis, autosomal recessive, with pilomatricomas |
| 12 | BMPRA1 | Juvenile polyposis |
| 13 | SMAD4 |  |
| 14 | VHL | Von Hippel–Lindau syndrome |
| 15 | MEN1 | Multiple endocrine neoplasia type 1 |
| 16 | RET | Multiple endocrine neoplasia type 2 & Familial medullary thyroid cancer |
| 17 | PTEN | PTEN hamartoma tumor syndrome |
| 18 | RB1 | Retinoblastoma |
| 19 | SDHD | Hereditary paraganglioma-pheochromocytoma syndrome |
| 20 | SDHAF2 |  |
| 21 | SDHC |  |
| 22 | SDHB |  |
| 23 | MAX |  |
| 24 | TMEM127 |  |
| 25 | TSC1 | Tuberous sclerosis complex |
| 26 | TSC2 |  |
| 27 | WT1 | WT1-related Wilms tumor |
| 28 | NF2 | Neurofibromatosis type 2 |
| 29 | COL3A1 | Ehlers-Danlos syndrome, vascular type |
| 30 | FBN1 | Marfan syndrome, Loeys-Dietz syndromes, and familial thoracic aortic aneurysms and dissections |
| 31 | TGFBR1 |  |
| 32 | TGFBR2 |  |
| 33 | SMAD3 |  |
| 34 | ACTA2 |  |
| 35 | MYH11 |  |
| 36 | MYBPC3 | Hypertrophic cardiomyopathy, dilated cardiomyopathy |
| 37 | MYH7 |  |
| 38 | TNNT2 |  |
| 39 | TNNI3 |  |
| 40 | TPM1 |  |
| 41 | MYL3 |  |
| 42 | ACTC1 |  |
| 43 | PRKAG2 |  |
| 44 | GLA |  |
| 45 | MYL2 |  |
| 46 | LMNA |  |
| 47 | FLNC |  |
| 48 | TTN |  |
| 49 | RYR2 | Catecholaminergic polymorphic ventricular tachycardia |
| 50 | CASQ2 |  |
| 51 | TRDN |  |
| 52 | PKP2 | Arrhythmogenic right ventricular cardiomyopathy |
| 53 | DSP |  |
| 54 | DSC2 |  |
| 55 | TMEM43 |  |
| 56 | DSG2 |  |
| 57 | KCNQ1 | Romano-Ward long-QT syndrome types 1, 2, and 3, Brugada syndrome |
| 58 | KCNH2 |  |
| 59 | SCN5A |  |
| 60 | LDLR | Familial hypercholesterolemia |
| 61 | APOB |  |
| 62 | PCSK9 |  |
| 63 | ATP7B | Wilson disease |
| 64 | OTC | Ornithine transcarbamylase deficiency |
| 65 | BTD | Biotinidase deficiency |
| 66 | GAA | Pompe Disease |
| 67 | RYR1 | Malignant hyperthermia susceptibility |
| 68 | CACNA1S |  |
| 59 | HFE | Hereditary hemochromatosis |
| 70 | ACVRL1 | Hereditary hemorrhagic telangiectasia |
| 71 | ENG |  |
| 72 | HNF1A | Maturity-onset diabetes of the young |
| 73 | RPE65 | RPE65-related retinopathy |

**Table 2.** PAS-GDC database description and statistics. PAS-GDC database includes genes, diseases, International Classifications of Disease (ICD) codes 9 and 10, as well as the relevant gene-disease combinations for each ICD code.

| Categories | Count |
| --- | --- |
| Genes | 73 |
| Diseases | 1788 |
| ICD 9 | 2101 |
| ICD 10 | 2589 |
| Gene-Disease Combination (ICD 9) | 7918 |
| Gene-Disease Combination (ICD 10) | 11,799 |

**Figure 1.**
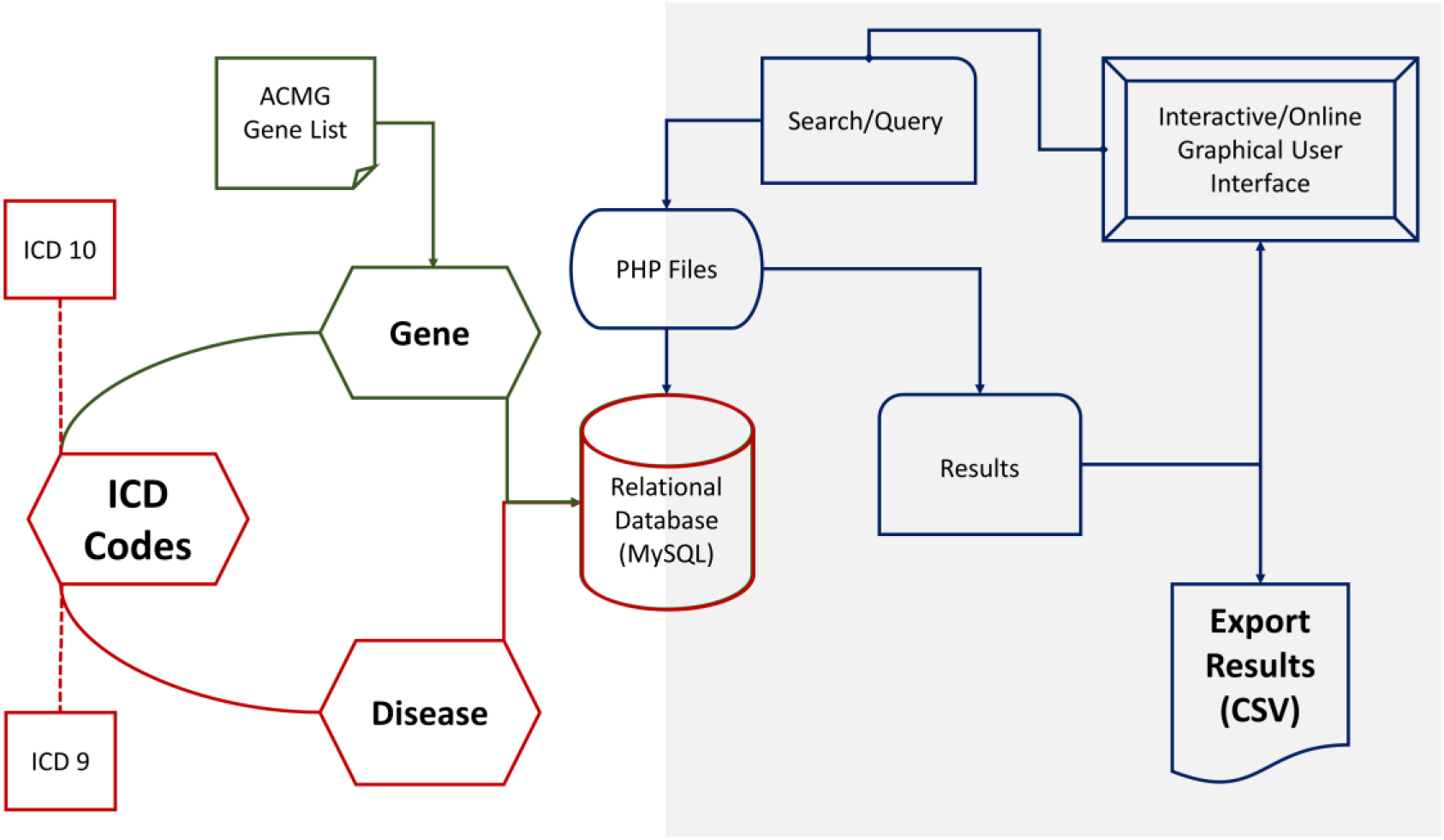
PAS: GDC components design, development, and data flow. PAS-GDC is an online application developed using MySQL database, PHP scripting language, and UNIX-based web and database servers.

### 2.2. Relational Database Modelling

The main objective of the database was to make the compiled information easily searchable and parsed so that all searches from the website would be up to date. Additionally, the database design needed to support easy integration of future ICD codes to ensure up-to-date information is reflected on our website. To meet these requirements, the database was created in MySQL Workbench and consisted of seven relations. The seven relations included ACMG’s 73 actionable genes, diseases, ICD 9 codes, ICD 10 codes, gene-disease pairings, gene-disease-ICD 9 pairings, and gene-disease-ICD 10 pairings. The gene-disease-ICD 9 and gene-disease-ICD 10 relations are created from the relations which manage the genes, diseases, and respective ICD codes (Figure 2). This ensures that there are no duplicate values. This database is unique and accessible through our freely available, open-source web application.

**Figure 2.**
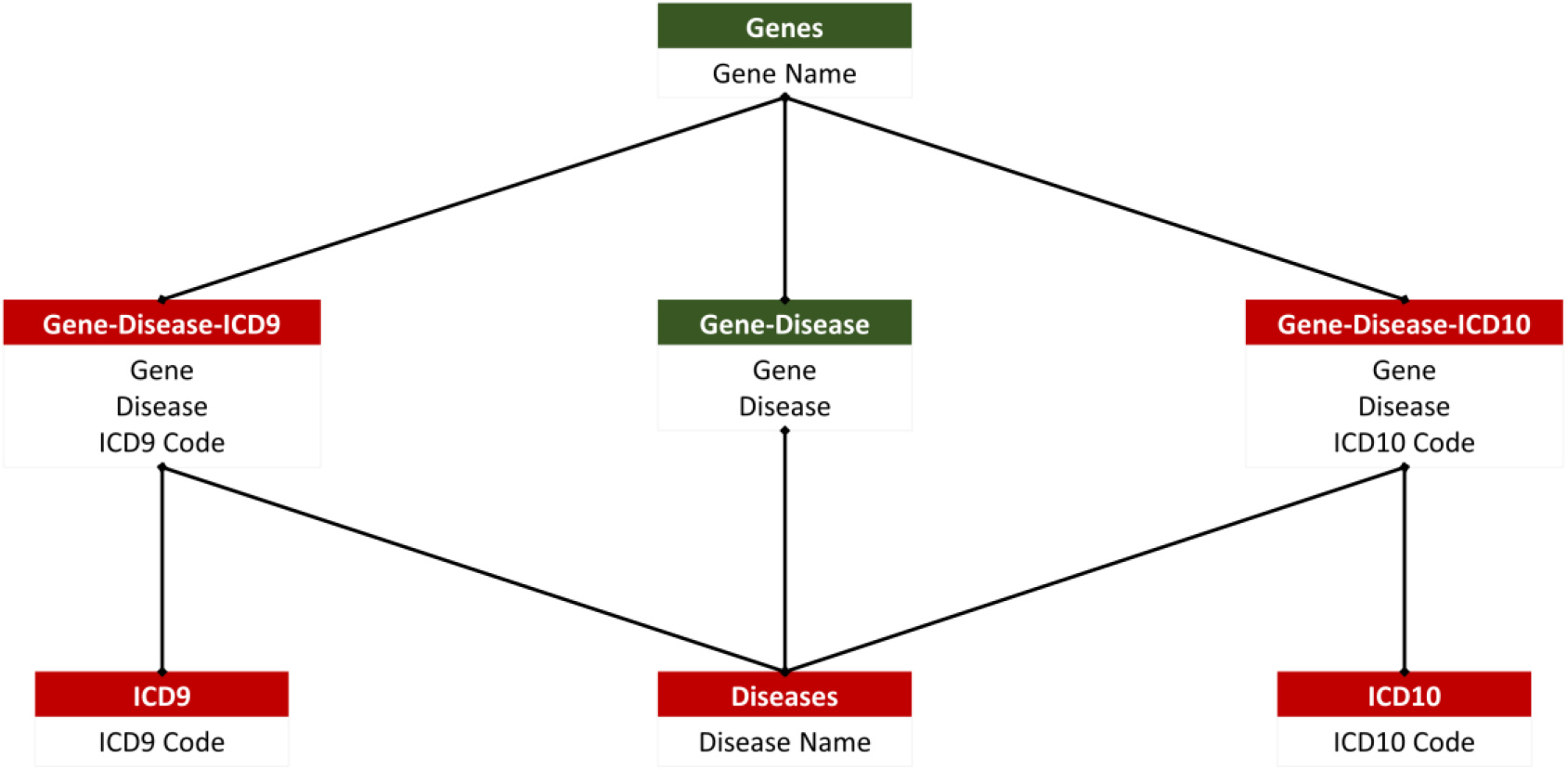
PAS: GDC Relational database. PAS-GDC database includes six relations, genes, diseases, ICD9, ICD10, gene-disease, gene-disease-icd9, and gene-disease-icd10.

### 2.3. Web development and search

PAS-GDC is a web application that has been developed using Hypertext Markup Language (HTML) and JavaScript with its jQuery packages. Additionally, we have used Cascading Style Sheets (CSS) with a Bootstrap framework on HTML to enhance the presentation and provide a user-friendly interface to our users. The database was connected to our web application using server-side PHP language and its ‘mysqli’ packages. Visual Studio Code was the primary Integrated Development Environment (IDE) used in the creation of the source code as well as testing. The testing of the website involved using RedHat localhost servers. During development, testers used macOS, iOS, Windows, and Android operating systems along with a variety of different browsers that include but are not limited to Google Chrome, Safari, and Firefox to ensure that the website performs typically and is configured correctly regardless of the environment. SSL certificates were utilized in the PAS-GDC website and the communication between the browser and server was encrypted. The search allows for the user to perform searches based on the ICD-9 codes, ICD-10 codes, Gene, or Disease category (Figure 3). The ICD-9 and ICD-10 searches allow for their independent gene and disease search allowing the user to retrieve the respective gene-disease pairing based on the desired ICD selection. Additionally, the website allows for a simple and easy export feature that allows the user to store and share their desired result as a CSV file.

**Figure 3.**
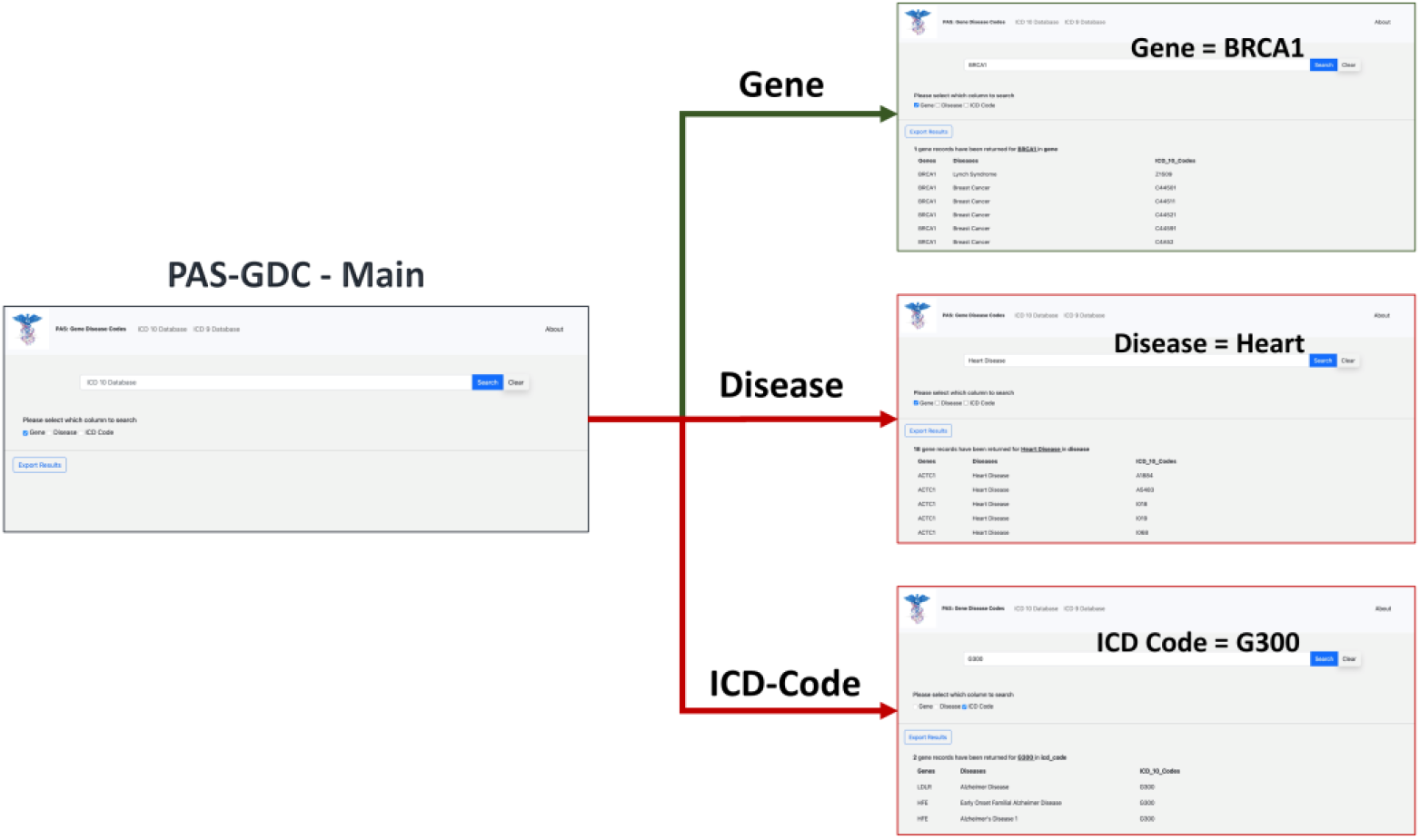
PAS: GDC graphical use interfaces (GUI) workflow. PAS-GDC GUI includes, Main, Gene, Disease, and ICD-Code (9 and 10) interfaces.

## 3. Results

The gene-disease-ICD code database is a flexible and dynamic database. The database design in SQL allows for more genes and ICD codes to be integrated as they are made available as well as it to be updated automatically on the PAS-GDC website. PAS-GDC is a simple-to-use, robust search engine that utilizes minimalistic features and an internet connection to retrieve results. The Graphical User Interface (GUI) includes a search capability of three features, namely (1) ICD Codes, (2) Genes, (3) Diseases as a simple check box that gives the user the capability to choose the feature they desire. Additionally, PAS-GDC provides the users the option to search their results against ICD-9 codes and ICD-10 codes.

### 3.1. Case-study: Gene

To test the effectiveness and functionality of the PAS-GDC web application, we created three different case studies exploring the ‘gene’ search feature (Figure 4). The genes that were included in this case-study were *BRCA1, MYBPC3*, and *APC*. The results were exported and collected in a tabular format with three columns: genes, diseases, and ICD-codes. The *BRCA1* gene codes for proteins that are vital to a multitude of cellular processes [30]. Mutations in this gene can lead to a predisposition of breast and ovarian cancers [30, 31, 32]. The search results for the *BRCA1* gene present fifty-seven distinct diseases that are directly linked to this gene. These diseases include but are not limited to breast, ovarian, and pancreatic cancer as well as fallopian tube carcinoma. Additionally, the search uncovered a total of 126 ICD-9 (Figure 4. A1) and 243 ICD-10 codes associated with *BRCA1* (Figure 4. A2). ICD-9 codes starting with 17 seemed to repeat for the BRCA1 gene as this category denotes breast cancer. Similarly, ICD-10 codes staring with C4 and C5 were most common in the search. The search criteria were repeated for the gene *MYBPC3*. Mutations in this gene are usually linked to cardiovascular diseases like cardiomyopathy and atrial fibrillation [33]. Seventeen other diseases were also found to be linked to this gene through our web application. These diseases included but were not limited to diastolic heart failure, cardiac arrest, and heart disease. Currently, there are fifty-nine ICD-9 (Figure 4. B1) and 104 ICD-10 codes linked to *MYBPC3* (Figure 4. B2). One of the most common diagnoses linked to this gene was cardiovascular diseases (heart disease), the leading cause of death in the United States [34, 35]. Thirty-two of the ICD-9 codes and fifty-seven of the ICD-10 codes are linked to heart disease showing its prevalence and impact on the patients with a genetic mutation in the *MYBPC3* gene. The third case study focused on the APC gene which is known to lead to a predisposition to colorectal and lung cancer [36, 37]. Based on results from the PAS-GDC web application, it was observed that there are forty-three diseases that are associated with this gene. Some of the diseases highlighted in the results included but were not limited to lung cancer susceptibility, thyroid, and breast cancer. Seventy-six ICD-9 (Figure 4. C1) and 186 ICD-10 codes were retrieved for the *APC* gene (Figure 4. C2). Notably, the most common diagnoses linked to this gene included breast and lung susceptibility cancer. While APC had been linked to lung cancer in previous studies, the relation with breast cancer has not yet been established. fifteen ICD-9 and seventy-nine ICD-10 codes for the APC gene were associated with breast cancer.

**Figure 4.**
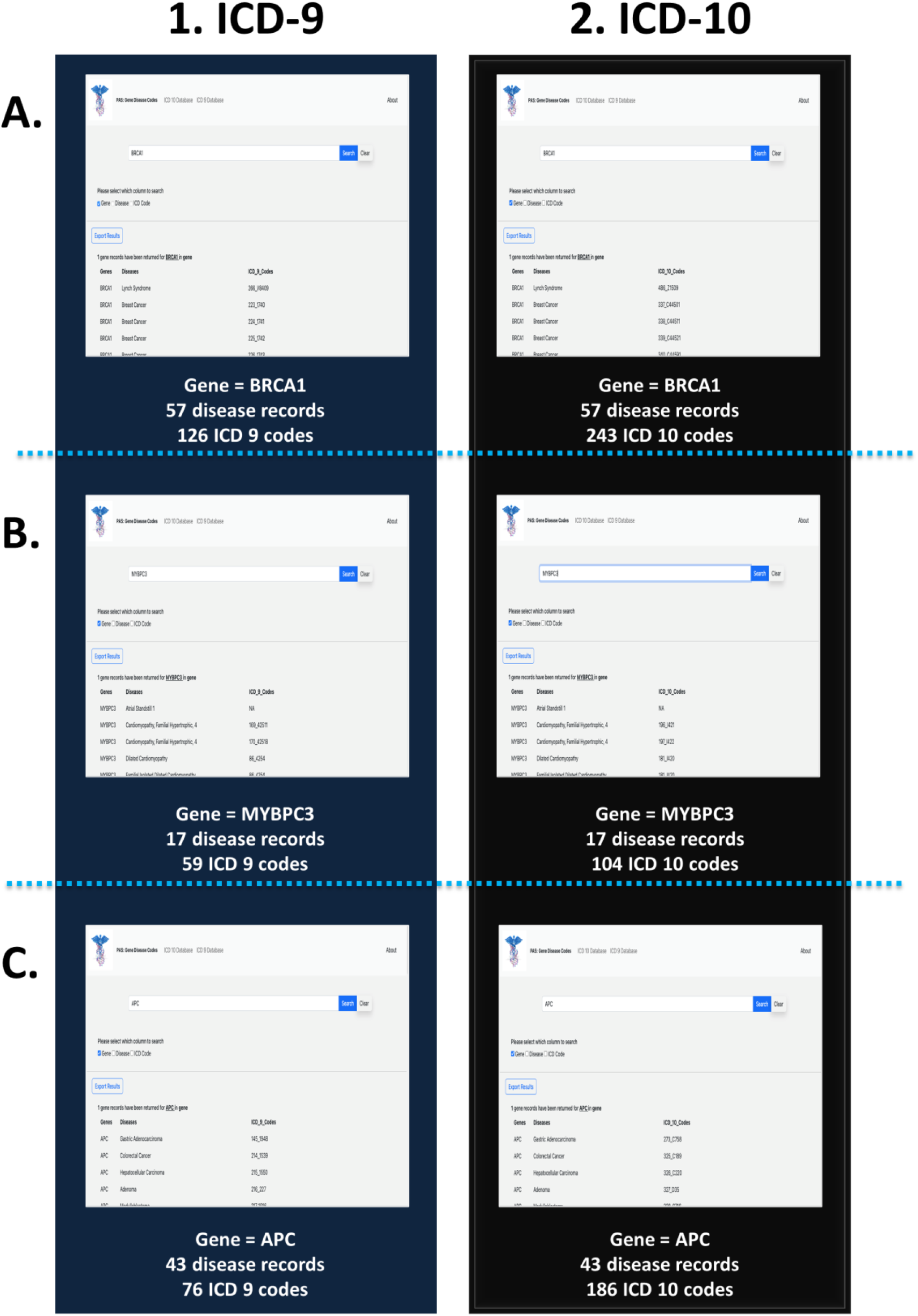
PAS: GDC use case - gene. This figure presents three different case studies exploring the ‘gene’ search feature: BRCA1, MYBPC3, and APC.

### 3.2. Case-study: Disease

The link between common disease nomenclature and international classification was exemplified through three case studies of breast cancer, heart disease and Alzheimer’s disease (Figure 5). The disease case studies were chosen because of their prevalence in the general population and the demonstrated interest in various fields by way of new research, and public and private funding. Breast cancer is one of the leading causes of death for women worldwide has an incidence of one in ten cancer diagnoses each year [38, 39]. Our web application highlights the impact of this disease by returning 323 ICD 9 (Figure 5. A1) and 1449 ICD 10 codes (Figure 5. A2). Additionally, a total of sixteen gene records were retrieved for breast cancer which includes but is not limited to *BRCA1, RB1, APC*, and *PTEN*. Like cancer, the term ‘heart disease’ encompasses several different subtypes, and one of the most common forms, congenital heart disease, continues to be a growing burden on healthcare systems [40]. A search of heart disease retrieved 540 ICD-9 (Figure 5. B1) and 937 ICD-10 codes (Figure 5. B2). A total of eighteen gene records were retrieved for heart disease and the most common gene being ACTC1 which has been documented to cause cardiomyopathies [41]. Our final case of Alzheimer’s is a disease that is characteristically prevalent in older adults and current research implicates a complex relationship between genetic and environmental factors [42]. The search yielded three ICD-9 (Figure 5. C1) and nine ICD-10 codes (Figure 5. C2). Additionally, two gene records, *LDLR* and *HFE*, were associated with this disease. While these genes have been studied in other forms of neurological diseases, there effects and interactions on neurodegenerative diseases are not as widely studied. Since ICD-10 diagnostic code set allows for greater specificity in the disease etiology, anatomic site, and severity [43], there is a greater number of codes available, as seen in all three case studies.

**Figure 5.**
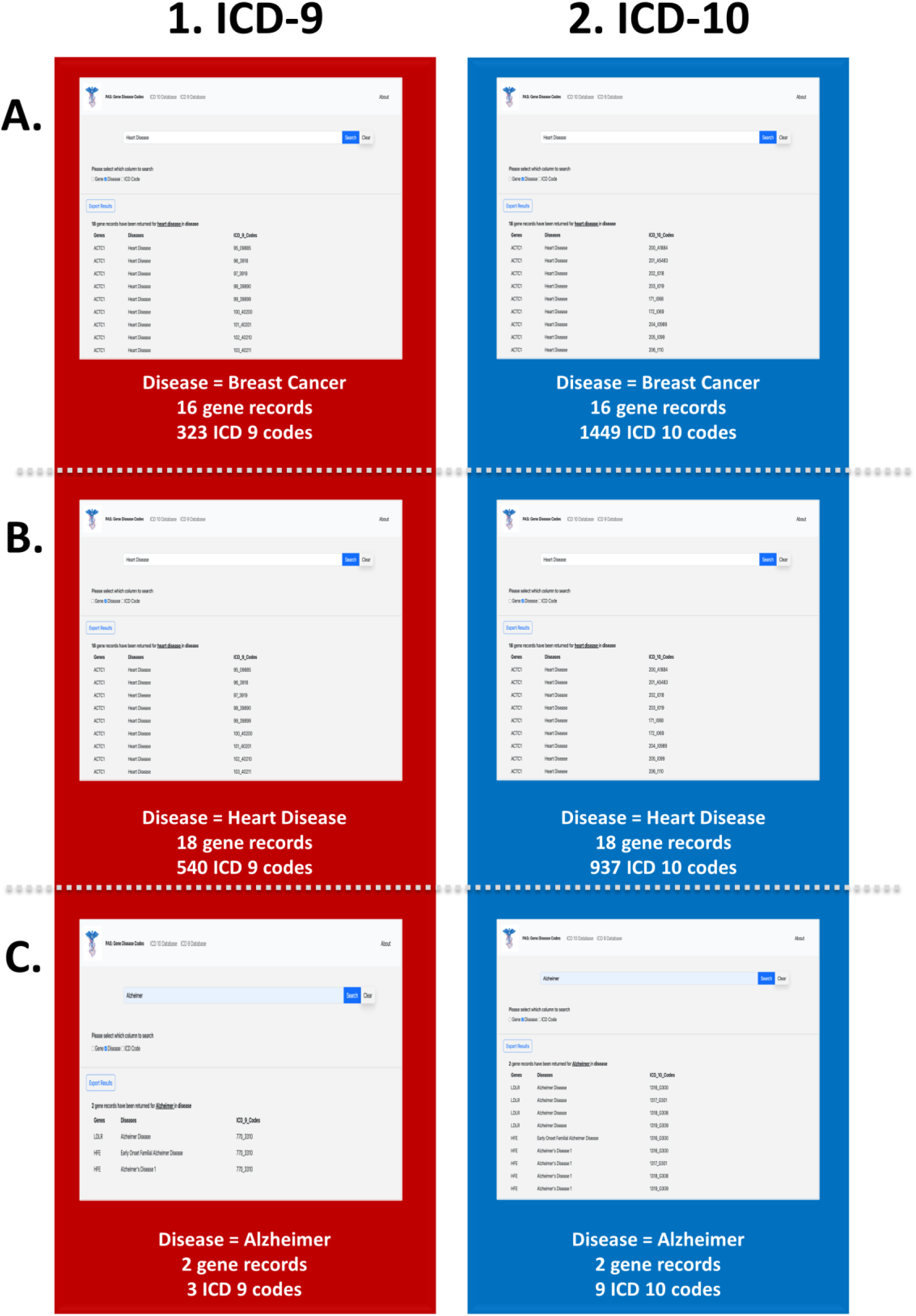
PAS: GDC use case - disease. This figure presents three different case studies exploring the ‘disease’ search feature: breast cancer, heart disease and Alzheimer’s disease.

### 3.3. Case-study: ICD - Code

The third search feature utilized by our web application is based on the ICD codes. The three ICD 9 codes that were included in this case-study were 331, 220, and 770. A search on our web application for the ICD 9 code 331 returns two unique genes, *APOB* and *LDLR*, as well as one common disease, vascular disease (Figure 6. A1). Notably, the LDLR gene is also associated with other forms of heart disease as stated previously. The ICD 9 code 233 yields one distinct disease, breast cancer, which is most common type of cancer (Figure 6. B1). Additionally, the search returns sixteen unique genes which includes but is not limited to *APC, BRCA1, MLH1, PTEN*, and *RB1*. A search of our third case-study for the ICD 9 code, 770, shows that this code is associated with Alzheimer’s as well as two distinct genes, *LDLR* and *HFE* (Figure 6. C1), which have been observed to be linked to Alzheimer’s based on previous queries. We also utilized our ICD 10 database for three distinct case studies involving the codes 411, 1316, and 202. A search of the ICD 10 code, 202, returns twelve unique genes and two diseases, heart disease and ptosis (Figure 6. A2). The code, 411, is associated mainly with breast cancer as well as other phenotypic variations of this disease such as breast giant fibroadenoma and breast benign neoplasm. Additionally, the search yields nineteen distinct genes which are linked to this unique ICD code (Figure 6. B2). The code 1316 is linked to one distinct disease, Alzheimer’s and two genes, *LDLR* and *HFE* (Figure 6. C2).

**Figure 6.**
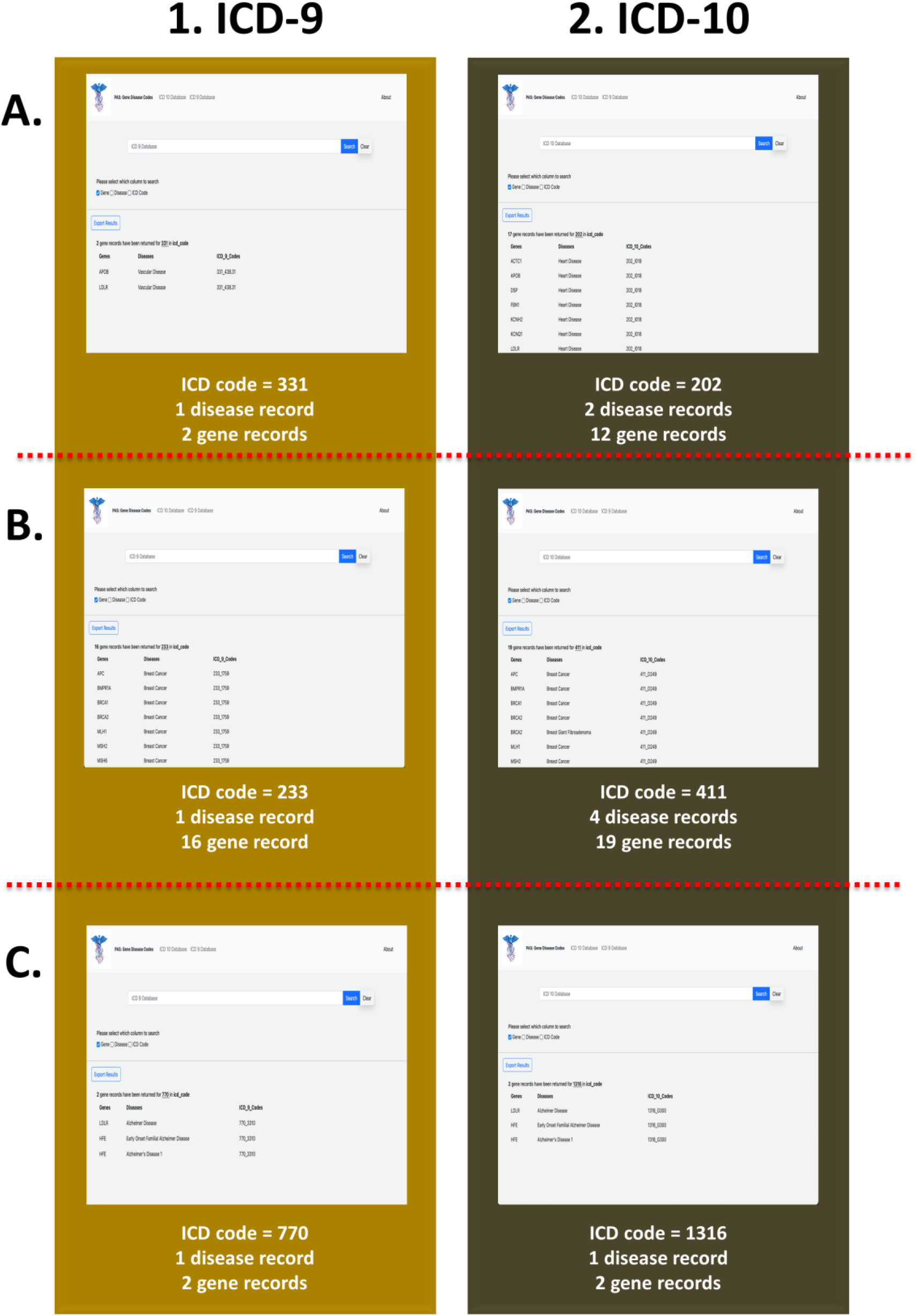
PAS: GDC use case – ICD code. Case-study: ICD - Code. This figure presents three different case studies exploring ICD-9 and ICD-10 codes.

The connection to our inhouse database does not require the user to install any tools or external modifications. When the user searches their desired keyword, it triggers the database and cross-references for exact or similar keywords. Once the database retrieves the results, it is presented to the user in a table format displaying the gene, disease, and ICD code as separate columns. Additionally, our web application allows the user to save their desired results as a text (CSV) file. The intelligent search feature of the PAS-GDC removes the need to cross verify genes or diseases on other web applications or databases by integrating and providing an all in one (Gene, ICD, Disease) search capability to the user. Updates to the database might include but are not limited to new additions to the ACMG genes, new associations between genes and diseases, and the addition of another version of the ICD code.

## 4. Discussion

Recent developments in the sequencing technologies and analysis of gene expression and variant data have helped advance the field of precision medicine [37]. Beyond isolated, single-based mutations, genomic and transcriptomic analyses have the potential to be a driver of clinically reliable predictions of complex disease and disorders. In a clinical research setting, the exploratory and dynamic nature of precision medicine yields promising results in discovering new gene-disease relationships, variants, and diverse genotyping [3]. These ideas are marketed by popular genomic analytic companies to the average consumer, but without the rigorous academic regulation and scrutiny. NGS has aided in the implementation of personalized treatments for patients with cardiovascular disease and neurodegenerative conditions [3]. Some of the applications of NGS include but are not limited genomic data models to support clinical decision making, identification of robust epigenic biomarkers as well as clinical translation [3]. Additionally, the latest research indicates that there is merit in integrating untargeted metabolomic profiling with genomic analysis for individuals at the ends of phenotypic expression [45]. This approach demonstrates that integrated genomics helps narrow the gap between treatment and disease by leveraging streamlined analysis on a patient’s genome. Thus, saving critical diagnosis time and money for the patient and institution of care [46]. However, there are still many constraints when trying to integrate genomic and clinical data. These constraints include but are not limited to lack of standardization when linking genes to their disease phenotypes [20], difficulty in integrating huge amounts of genetic and clinical data [47] and absence of a single application platform that contains up-to-date genome and clinical data [20, 48, 49]. To address these limitations, we have created PAS-GDC, a web application that is easy to navigate, and freely available on many platforms. This GUI was designed so that it can be used by non-computational users, such as physicians and geneticists, allowing for the integration of precision medicine in the clinical field.

One of the immediate implications of our web application is the downstream bioinformatic analysis involving gene-disease relationships. We have previously proposed a similar model called GVViZ, a Findable, Accessible, Interactive, and Reusable (FAIR) application that is available across platforms for RNA sequence-driven variable and complex gene-disease annotations and expression through dynamic heat map visualizations [50, 51]. Beyond a computational lens, clinicians and patients can interpret clinical and genomic data by learning the implications of one or more mutations in their genome. From the clinician’s perspective, they could present actionable steps in a more effective, personalized treatment plan. Additionally, researchers in various fields could use our web application to support their work, especially those seeking connections between genomics and a phenotypical manifestation.

One of the main limitations of our project is the manual curation of the database. The fundamental aspect of our database is the 73 ACMG codes as well as the ICD-9 and ICD-10 codes. The backend development was labor intensive, and we chose actionable genes that have been shown to be causative of disorders as a strategic start. There are seven relationship databases created to compile the information cohesively and serves as a base for future updates, allowing our web application to remain up to date. To optimize our process, we are exploring different methods to address the time-consuming aspect of data curation. Furthermore, we are interested in using machine learning (ML) and Artificial Intelligence (AI) algorithms for data mining. A solid foundation was created and the tools to build out the data base to a more robust capacity are readily available. We are extending the scope of our project by implementing more disease-causing genes in our database as well as different versions of the ICD code as they are made available. With the copious amounts of data available, and the development of systems that can interpret them on a large scale, the focus of treatment can shift from symptom-based to prevention and early intervention in unprecedented ways. A world with precision medicine would challenge the current healthcare system by centering care around maintaining health instead of addressing the lack thereof.

## List of Abbreviations

(ACMG): American College of Medical Genetics and Genomics
(ASCII): American Standard Code for Information Interchange
(CSS): Cascading Style Sheets
(DNA): Deoxyribose Nucleic Acid
(EHR): Electronic Health Records
(FDA): Food and Drug Administration
(FAIR): Findable, Accessible, Interactive, and Reusable
(GVViZ): Visualizing Genes with disease causing Variants
(GUI): Graphical User Interface
(HTML): Hypertext Markup Language
(ICD): International Classifications of Disease
(IDE): Integrated Development Environment
(NGS): Next Generation Sequencing
(NDC): National Drug Code
(PAS): PROMIS-APP-SUITE
(GDC): Gene Disease Code
(SAM): Sequence Alignment Map
(SQL): Structured Query Language
(VCF): Variant Call Format
(WHO): World Health Organization
(WGS): Whole Genome Sequencing
(WES): Whole Exome Sequencing

## Acknowledgements

We appreciate great support by the Rutgers Institute for Health, Health Care Policy, and Aging Research (IFH); Department of Medicine, Rutgers Robert Wood Johnson Medical School (RWJMS); and Rutgers Biomedical and Health Sciences (RBHS), at the Rutgers, The State University of New Jersey. We thank members and collaborators of Ahmed Lab at the Rutgers (IFH, RWJMS, RBHS) for their support, participation, and contribution to this study.

This study was completed in part by research services and/or survey/data resources provided by the Institute for Health Survey / Data Core at Rutgers University.

The authors acknowledge the Office of Advanced Research Computing (OARC) at Rutgers, The State University of New Jersey for providing access to the Amarel cluster and associated research computing resources that have contributed to the results reported here.

## Author contributions

Z.A. proposed, supervised, and led this study. R.W. programmed the online/web interface of the application. A.P. modelled relational database, and implemented data extraction, transfer, and loading (ETL) modules to efficiently parse and insert data into designed database. K.P. and A.S.N. performed in data curation, integration, and management. W.P.L. and H.A. participated in post development analysis, and theoretical research. H.A., S.B. and S.M. did quality testing. All authors have participated in writing and review, and have approved manuscript for publication.

## Author information

R.W. is the Research Assistants at the Ahmed lab, Rutgers Institute for Health, Health Care Policy and Aging Research, Rutgers University-New Brunswick.

A.P. is the Research Assistants at the Ahmed lab, Rutgers Institute for Health, Health Care Policy and Aging Research, Rutgers University-New Brunswick.

A.S.R. is the Research Assistants at the Ahmed lab, Rutgers Institute for Health, Health Care Policy and Aging Research, Rutgers University-New Brunswick.

W.P.L. is the Research Assistants at the Ahmed lab, Rutgers Institute for Health, Health Care Policy and Aging Research, Rutgers University-New Brunswick.

S.B. is an intern at the Ahmed lab, Rutgers Institute for Health, Health Care Policy and Aging Research, Rutgers University-New Brunswick.

S.M. is an intern at the Ahmed lab, Rutgers Institute for Health, Health Care Policy and Aging Research, Rutgers University-New Brunswick.

K.P. was the Research Assistants at the Ahmed lab, Department of Genetics and Genome Sciences, UConn Health.

H.A. is the senior Research Assistants at the Ahmed lab, Rutgers Institute for Health, Health Care Policy and Aging Research, Rutgers University-New Brunswick.

Z.A. is the Assistant Professor of Medicine – Tenure Track and Core Member at the Rutgers Institute for Health, Health Care Policy and Aging Research; and Department of Medicine – Division of General Internal Medicine, Rutgers Robert Wood Johnson Medical School, Rutgers Biomedical and Health Sciences, Rutgers University-New Brunswick. ZA is the Adjunct Assistant Professor at the Department of Genetics and Genome Sciences, UConn School of Medicine, UConn Health, CT; and Full Academic Member of the Rutgers Microbiology and Molecular Genetics; Center for Cancer Health Equity, Rutgers Cancer Institute of New Jersey; Rutgers Human Genetics Institute of New Jersey, NJ.

## Declarations

### Ethical Approval and Consent to participate

Not applicable.

### Consent for publication

Not applicable

### Availability of data and material

The datasets used and analyzed during the current study are freely accessible through website: < https://promis.rutgers.edu/pas/>

### Data Availability Statement

The data used in the current study are available from the corresponding author on reasonable request.

## Availability and requirements

**Project name:** PAS-GDC

**Operating system:** Cross platform (Microsoft Windows, MAC, Unix, Linux)

**Programming languages:** HTML/CSS with Bootstrap Framework, PHP, MySQL, JavaScript

**Requirements:** The developer is responsible for MySQL installation and database schema.

**License:** Freely distributed for global users. Any restrictions to use by non-academics: Copyrights are to the authors.

**Download link: PAS: GDC** freely available and can be accessible through <https://promis.rutgers.edu/pas/>.

